# Torix group *Rickettsia* are widespread in New Zealand freshwater amphipods: using blocking primers to rescue host COI sequences

**DOI:** 10.1101/2020.05.28.120196

**Authors:** Eunji Park, Robert Poulin

## Abstract

Endosymbionts and intracellular parasites are common in arthropods and other invertebrate hosts. As a consequence, (co)amplification of untargeted bacterial sequences has been occasionally reported as a common problem in DNA barcoding. The bacterial genus *Rickettsia* belongs to the order Rickettsiales and consists of two lineages: one including diverse pathogens infecting arthropod hosts, the other consisting of non-pathogenic species with a broader host taxonomic range. While discriminating among amphipod species with universal primers for the COI region, we unexpectedly detected rickettsial endosymbionts belonging to the Torix group. To map the distribution and diversity of *Rickettsia* among amphipods hosts, we conducted a nationwide molecular screening of seven families of freshwater amphipods collected throughout New Zealand. In addition to uncovering a diversity of Torix group *Rickettsia* across multiple amphipod populations from three different families, our research indicates that 1) detecting Torix *Rickettsia* with universal primers is not uncommon, 2) obtaining ‘*Rickettsia* COI sequences’ from many host individuals is highly likely when a population is infected, and 3) obtaining ‘host COI’ may not be possible with a conventional PCR if an individual is infected. Because *Rickettsia* COI is highly conserved across diverse host taxa, we were able to design blocking primers that can be used in a wide range of host species infected with Torix *Rickettsia*. We propose the use of blocking primers to circumvent problems caused by unwanted amplification of *Rickettsia* and to obtain targeted host COI sequences for DNA barcoding, population genetics, and phylogeographic studies.

## Introduction

The cytochrome c oxidase subunit 1 gene (COI), a partial fragment of mitochondrial DNA, is the marker of choice for DNA barcoding, and is also widely used for population genetics and phylogeographic studies (Bucklin et al., 2011; Hajibabaei et al., 2007; Hebert et al., 2004). A variable region is flanked by highly conserved regions; this allowed for the design of a pair of universal primers and their application to various organisms (Folmer et al., 1994; Hebert et al., 2003). With the advancement of fast and cost-effective next-generation sequencing technologies, which enables metabarcoding (Elbrecht and Leese, 2015; Taberlet et al., 2012), the number of COI sequences is increasing rapidly in public databases such as GenBank and The Barcode of Life DataSystems (BOLD) (Porter and Hajibabaei, 2018). However, quality control is often an issue due to the presence of questionable “COI-like” sequences (Buhay, 2009) or nuclear mitochondrial pseudogenes (numts) that are often coamplified with orthologous mtDNA (Song et al., 2008). Bacterial sequences are also often coamplified with universal primers. Indeed, there have been reports of the amplification of untargeted sequences of endosymbiotic bacteria such as *Wolbachia* and *Aeromonas* during DNA barcoding with universal primers and their misidentification as those of invertebrate hosts during deposition in databases (Mioduchowska et al., 2018; Smith et al., 2012).

The bacterial genus *Rickettsia* is another of these endosymbiotic taxa. This genus belongs to the order Rickettsiales along with *Wolbachia*, and comprises diverse pathogenic species that cause vector-borne diseases in birds and mammals including humans. Some rickettsioses (diseases that are caused by *Rickettsia*) with severe symptoms are well known, and include Rocky Mountain spotted fever, Queensland tick typhus, Rickettsial pox, and murine or epidemic typhus (Parola et al., 2005; Perlman et al., 2006; Weinert, 2015). To date, at least 13 groups are known within the genus *Rickettsia*: Adalia, Bellii, Canadensis, Guiana, Helvetica, Meloidae, Mendelii, Rhyzobious, Spotted fever, Scapularis, Torix, Transitional, and Typhus (Binetruy et al., 2020; Hajduskova et al., 2016; Weinert et al., 2009). All these groups except the Torix group are exclusively associated with arthropod hosts, such as mites, fleas, ticks, and spiders. The Torix group, which is sister to all other groups, is the only group that includes non-arthropod hosts such as amoeba and leeches (Galindo et al., 2019; Kikuchi et al., 2002; Kikuchi and Fukatsu, 2005). In addition to these freshwater hosts, the Torix group occurs in diverse arthropod groups that spend part of their life cycle in the aquatic environment (e.g. Coleoptera and Diptera) (Küchler et al., 2009; Pilgrim et al., 2017; Weinert et al., 2009).

Although *Rickettsia* are known as common pathogens or endosymbionts in arthropod hosts, this group has never been reported in crustaceans. *Rickettsia*-like organisms (RLO) have been reported in some groups of crustaceans including crabs, crayfish, lobsters, shrimps, and amphipods (Gollas-Galván et al., 2014). However, most reports of these RLOs were based on morphological similarity with *Rickettsia* and were rarely confirmed by molecular data. In amphipods, RLOs were reported in several species of gammarids, as well as other taxa (e.g., *Crangonyx floridanus* and *Diporeia* sp.) (Graf, 1984; Larsson, 1982; Messick et al., 2004; Winters et al., 2015). *16S rRNA* sequences of RLOs are available for *Diporeia* sp. and some gammarids, but none of them belong to the genus *Rickettsia* (Bojko et al., 2017; Winters et al., 2015).

In the course of an investigation of the diversity of New Zealand freshwater amphipods, we obtained a COI sequence that was highly divergent from that obtained from other populations (~57%) of the same host group (*Paracalliope* species complex). DNA from the individual amphipod was extracted from its legs (i.e., low chance of contamination due to gut contents). We obtained a clear chromatogram with no ambiguous peaks. Furthermore, this sequence was similar to other COI sequences obtained from diverse insects (Coleoptera, Diptera, Hemiptera, Hymenoptera, Odonata) in GenBank with sequence similarity ranging from 80 to 99 %. However, these sequences were also similar (~92%) to sequences of rickettsial endosymbionts of insects and spiders that have been recently registered in GenBank. Because such highly conserved COI sequences among distantly related arthropod groups are unlikely, we assumed that these sequences were actually obtained from their rickettsial endosymbionts. We independently confirmed the presence of *Rickettsia* in our amphipod hosts using three genetic markers that were designed to be specific to *Rickettsia*.

Because *Rickettsia* is an endosymbiont within host cells, DNA extracts from infected host tissue will inevitably include DNA of endosymbionts as well. If binding sites for ‘universal primers’ are conserved in both hosts and their endosymbionts, PCR products obtained from these mixed templates may result in mixed signals in chromatograms, or in the amplification of endosymbiont instead of host sequences. Using primers that are designed to bind uniquely and specifically to host templates would reduce this problem. However, designing group-specific primers is not always possible, especially when reference sequences are scarce or not available. Also, finding conserved regions across a given taxonomic group may not be achievable. Alternatively, blocking primers can be used to prevent the amplification of unwanted or dominant sequences among DNA templates (Vestheim and Jarman, 2008). For example, this method has been successfully applied to identify prey items (by suppressing the amplification of predator DNA in gut contents), or to obtain rare mammal sequences from ancient DNA (by blocking the amplification of human DNA) (Boessenkool et al., 2012; Vestheim et al., 2011). Because COI sequences of Torix *Rickettsia* are highly conserved in diverse host groups, we were able to design blocking primers that are intended to specifically block the amplification of Torix *Rickettsia* but allow amplification of the COI region of (any) host mtDNA.

In this study, we first screened rickettsial infections in diverse amphipods collected throughout New Zealand to determine their prevalence and distribution. Secondly, we characterized the genetic diversity of the newly found *Rickettsia* in relation to other Torix *Rickettsia* using 4 distinct markers, namely *16S rRNA*, *gltA*, *atpA*, and COI, and expand current understanding of *Rickettsia* phylogeny. Thirdly, we demonstrate that unwanted amplification of rickettsial COI sequences during host identification or barcoding is a common problem, and that such sequences have been frequently reported and misidentified in GenBank. Fourthly, we suggest that using blocking primers in addition to universal primers for PCR is an effective solution to obtain targeted host COI sequences. Finally, we discuss the implications of these pseudo-sequences in public databases, ways to reduce this problem, and possible applications of blocking primers for similar problems.

## Methods

### Confirming *Rickettsia* infections by PCR

The presence of *Rickettsia* was confirmed by amplification of three different markers (*16S rRNA*, *gltA*, and *atpA*) using *Rickettsia*-specific primer pairs (Küchler et al., 2009; Pilgrim et al., 2017) (Table 1). The DNA sample (S15_470) from which we obtained presumably ‘rickettsial COI’ was used as DNA templates. Seven additional DNA samples from the same population were also included for PCR detection to compare the efficiency of the primer sets in order to select the best marker for molecular screening. For PCR reactions, 12.3 μl of distilled water, 4 μl of reaction buffer, 0.8 μl of each forward and reverse primers, 0.1 μl of MyTaq (Bioline), and 2 μl of DNA were used. PCR reactions were conducted under the following conditions: 95°C initial denaturation for 5 min, 35 cycles of 94°C for 30s, 54°C for 30s, 72°C for 120s, final extension for 7 min at 72°C. Then, 2μl of PCR product from each PCR reaction was run on a 1.5 % agarose gel.

**Table 1.**
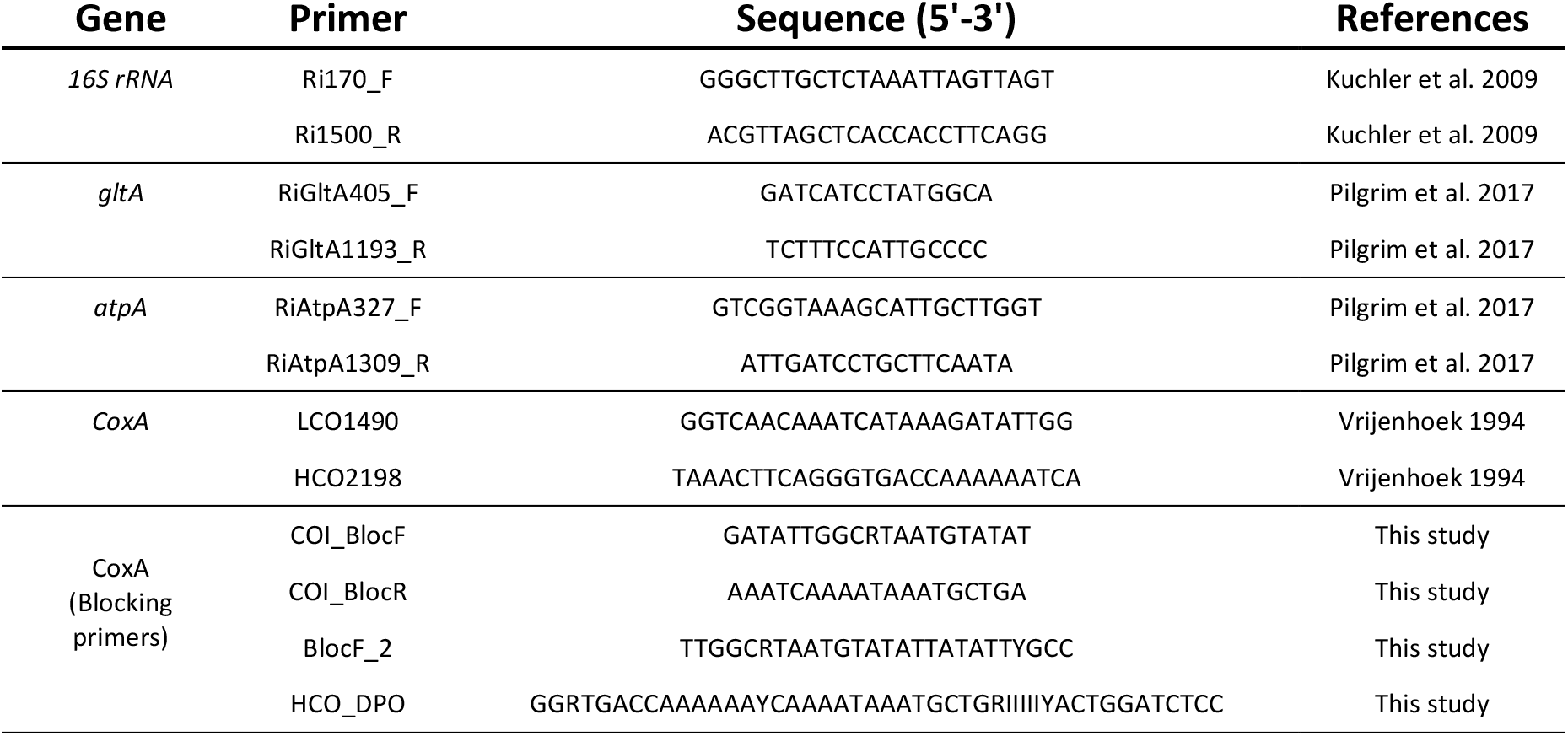
List of primers used in this study. All blocking primers were modified with C3 spacer at the 3’ end. Five deoxyinosines (I) were added in the middle of HCO_DPO primer as a linker.

### Molecular screening of *Rickettsia* in New Zealand amphipods

We obtained extracted DNA samples of diverse amphipod specimens from Park et al. (2020), in which the authors investigated the diversity of microsporidian parasites, a group of obligate intracellular eukaryotic parasites of amphipod hosts. Seven families of amphipods (Melitidae, Paracalliopidae, Paraleptamphopidae, Phreatogammaridae, Talitridae, Paracorophiidae, and an undescribed family of Senticaudata) were collected from 69 locations throughout both the South and North Islands (Figure 1 and Supplementary Table 1). A total of 724 pooled DNA samples obtained from 2,670 individuals (mostly 4 individual amphipods per pool) were screened for *Rickettsia* by PCR by amplifying the *16S rRNA* region under the PCR conditions and procedures described above.

**Figure 1.**
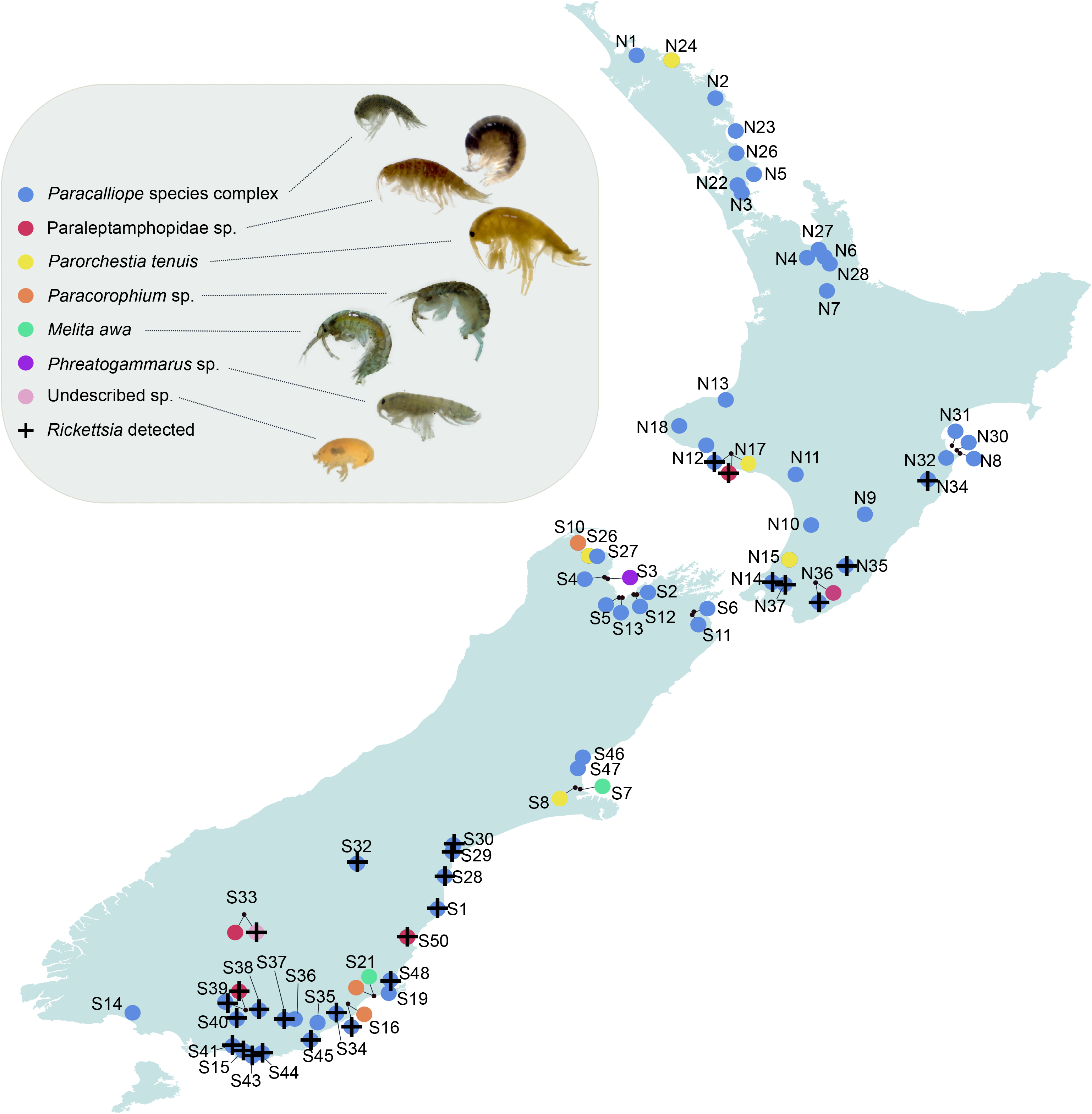
Map of sampling sites. Map of New Zealand showing the sixty-nine sampling sites with circles. Seven different families of amphipods are marked with different colours in the circles. The sites where *Rickettsia* was detected are marked with +. Site codes correspond to those in Supplementary Table1.

### Sequencing

*16S rRNA*, *gltA*, and *atpA* sequences were obtained from populations that had been found positive for *Rickettsia* infections by PCR screening (Supplementary Table 1). PCR products were purified with MEGAquick-spinTM Total Fragment DNA Purification Kit (iNtRON Biotechnology) following the manufacturer’s instructions. Purified PCR products were sent to Macrogen, Korea, for Sanger sequencing. Raw sequences were aligned in Geneious Prime 2019.0.4 (https://www.geneious.com) and ambiguous sites were carefully examined by eye. Haplotypes were identified by using the package *pegas* (Paradis, 2010) in R version 3.5.2 (R Development Core Team, 2011). We obtained some rickettsial COI sequences as a byproduct during host identification of amphipods with universal primers. We included these COI sequences along with nucleotide sequences from other genes for further analyses.

### BLAST search

Blast search was done on GenBank with the *16S rRNA*, *gltA*, *atpA*, and COI sequences obtained in this study. Based on the result of BLAST searches, all sequences that were considered as the Torix group (sequences with similarity to the query sequence higher than that between the query sequence and the Bellii group *Rickettsia*) were downloaded from GenBank (see Supplementary Tables 2-4) for further phylogenetic tree reconstructions. *16S rRNA* and *gltA* sequences from other *Rickettsia* groups were also obtained and included for tree reconstruction (Supplementary Table 5). In addition, *16S rRNA* sequences from recently discovered close relatives to *Rickettsia* (*Candidatus Trichorickettsia*, *Candidatus Gigarickettsia*, and *Candidatus Megaira*), and *Orientia tsutsugamushi* were included as outgroups.

### Phylogenetic analyses

For each gene set, all sequences were aligned in Geneious Prime with the MUSCLE algorithm. Ambiguous sites were then eliminated in Gblocks with the least restrictive setting (Castresana, 2000). The best-fitting model of nucleotide evolution for each dataset was determined based on the corrected Aikake information criterion (AICc) using jModelTest v2.1.6 (Darriba et al., 2012), which was conducted through the CIPRES Science Gateway v3.3 (Miller et al., 2010). For all datasets, the General Time Reversible (GTR) model of nucleotide substitution along with Gamma distributed rate variation across sites (G) and the proportion of invariable sites (I) were chosen as the best model. Bayesian trees were inferred in MrBayes 3.2.7a (Ronquist et al., 2012). For each dataset, two independent runs, which consisted of four chains each, were simultaneously conducted for 10,000,000 generations with a sampling frequency of 1000. The initial 25% of samples were discarded. The resulting trees were visualized in FigTree v1.4.4.

### Design of blocking primers

Blocking primers were designed following the guidelines of Vestheim et al. (2011) (Vestheim et al., 2011). In order to design blocking primers, COI sequences of the Torix group and some species belonging to other groups of *Rickettsia*, and COI sequences of New Zealand amphipods were aligned in Geneious Prime (Figure 2). We designed four different annealing inhibiting blocking oligos which were intended to compete with universal primers. All met the following criteria: First, the blocking primers should overlap with one of the universal primers. Second, the blocking primers should specifically bind to the unwanted DNA templates (i.e. *Rickettsia*) but not to our target DNA templates (i.e. amphipod hosts). Third, 3’-end was modified so that it does not prime amplification (here, all with C3 spacer). Initially, two primers were designed: Bloc_F and Bloc_R (Table 1). However, GC contents of these primers were too low (27.5% and 21%, respectively), which resulted in a low expected melting temperature (Tm) of 43.2°C and 42.5 °C, respectively. Ideally, Tm of a blocking primer should be higher than that of the competing primers (Vestheim et al., 2011). We, therefore, designed a longer primer, BlocF_2, to increase Tm to 51.7 °C (Table 1). A fourth primer was designed with a dual priming oligonucleotide (DPO): HCO_DPO (Table 1). A DPO can be used when it is impossible to find an appropriate binding site for a blocking primer adjacent to a binding site of a universal primer (Vestheim and Jarman, 2008). A DPO primer consists of two separate segments connected with five deoxyinosines, and with C3 spacer modification at the 3’end. The total length of a typical DPO primer is long but it does not suffer from high Tm because a deoxyinosine linker, which assumes a bubble-like structure, allows the two segments to act independently (Chun et al., 2007). All synthesized primers were purified with polyacrylamide gel purification (PAGE) to increase binding specificity by removing under-synthesized oligos.

**Figure 2.**
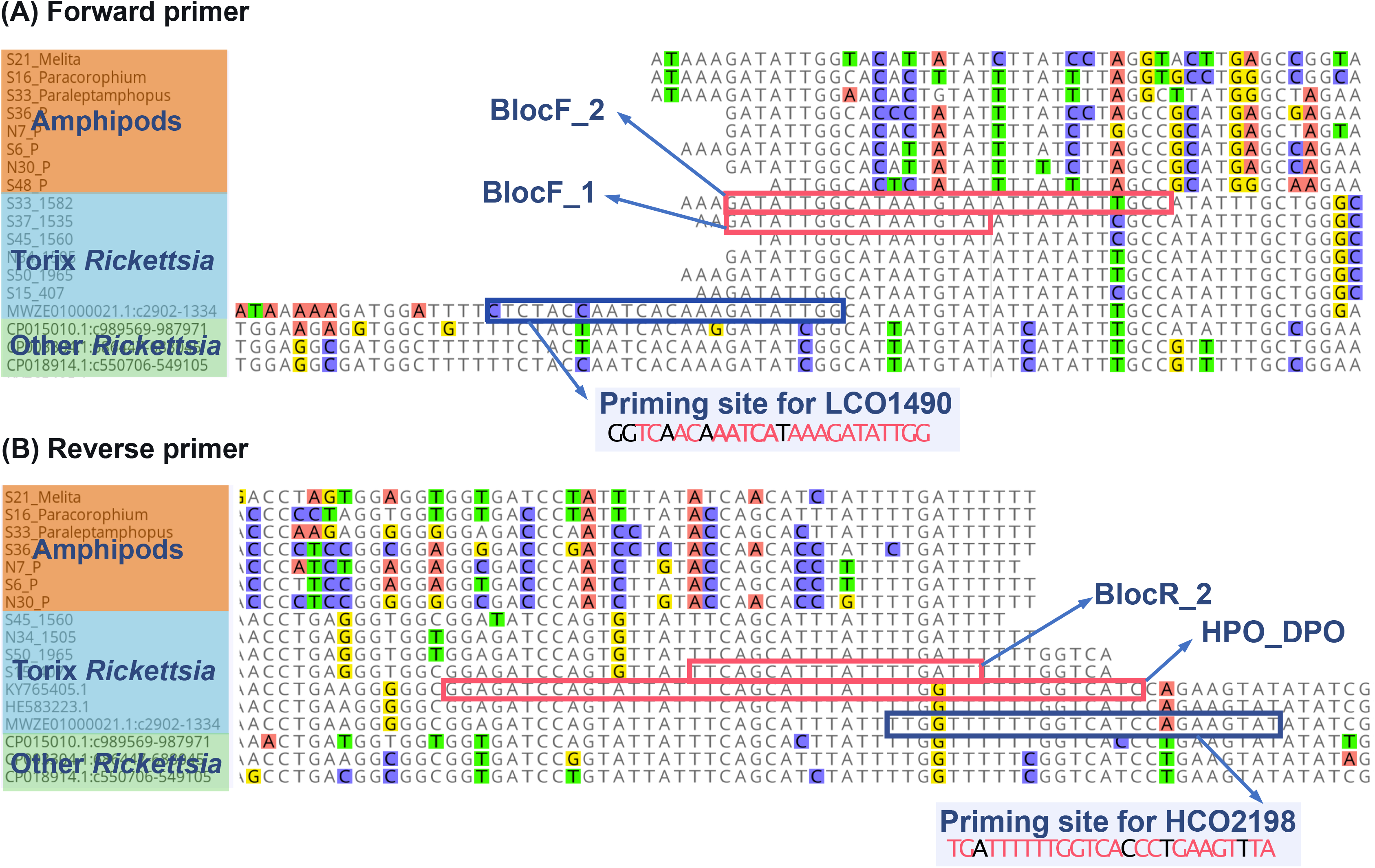
COI sequence alignments showing priming sites for both forward and reverse primers. Conserved regions to which universal primers bind are highlighted with blue squares. Positions at which nucleotides are the same as in universal primers are highlighted with pink texts in the primer sequences. COI sequences are highly divergent among amphipods whereas COI sequences from Torix *Rickettsia* from diverse host groups are highly conserved, which allowed the blocking primers (binding regions are highlighted with pink squares).

### Validation of blocking primers

We applied fragment analysis to test and compare the effectiveness of our blocking primers (Vestheim and Jarman, 2008). Fragment analysis of fluorescently labeled PCR products on capillary electrophoresis can separate fragments in different sizes and can be used as a semi-quantitative method. When amplified with the universal LCO1490 and HCO2198 primers, the expected lengths of PCR products were different for amphipod hosts and *Rickettsia* COI, because Rickettsial COI is 6 bp longer. The FAM dye was attached to the 5’ end of the LCO1490 primer. This fluorescently labeled forward primer was added to the PCR mixture instead of the normal (unlabeled) LCO1490 primer. Various factors can affect PCR success with blocking primers: Tm of primers, the concentration of primers (relative ratio between blocking primer and regular primer), the amount of the DNA templates in a PCR mixture (concentration of DNA), and the number of PCR cycles (Vestheim and Jarman, 2008). To optimize PCR conditions, PCR reactions were conducted under several different PCR conditions (Table 2). Fragment analyses were carried out with a 1,200 LIZ size marker on an ABI 3730xl System (Applied Biosystems) at Macrogen (Korea). Results were analyzed with Peak Scanner Software 1.0 (Applied Biosystems).

**Table 2.**
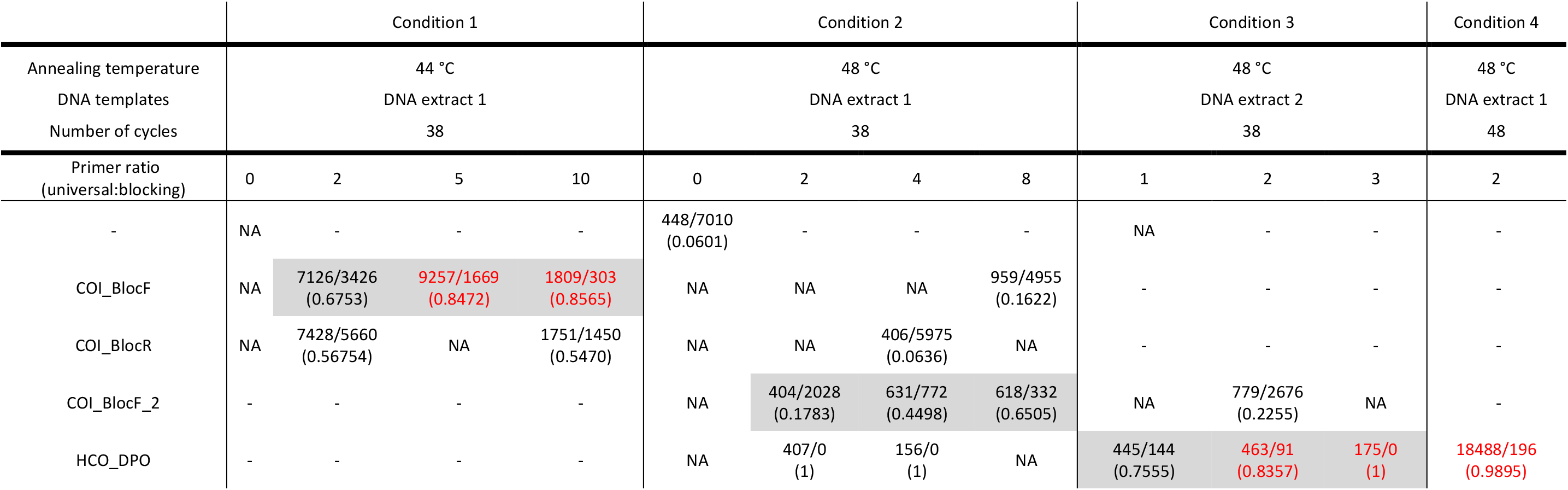
PCR conditions and results of fragment analyses. PCR products obtained under different PCR conditions (different primers with different concentrations, annealing temperature, different DNA templates, number of PCR cycles) were run on capillary electrophoresis. The number of amplicons of host COI is followed by the number of amplicons of *Rickettsia* COI separated by (/), and their ratio is shown in parenthesis. The effects of blocking primers are highlighted in the grey boxes. Universal to blocking primer ratios that were higher than 0.8 are highlighted in red. NA: fragment analysis was conducted but the result was not available.

|  | Condition 1 |  |  |  | Condition 2 |  |  |  | Condition 3 |  |  | Condition 4 |
| --- | --- | --- | --- | --- | --- | --- | --- | --- | --- | --- | --- | --- |
| Annealing temperature | 44 °C |  |  |  | 48 °C |  |  |  | 48 °C |  |  | 48 °C |
| DNA templates | DNA extract 1 |  |  |  | DNA extract 1 |  |  |  | DNA extract 2 |  |  | DNA extract 1 |
| Number of cycles | 38 |  |  |  | 38 |  |  |  | 38 |  |  | 48 |
| Primer ratio<br>(universal:blocking) | 0 | 2 | 5 | 10 | 0 | 2 | 4 | 8 | 1 | 2 | 3 | 2 |
| - | NA | - | - | - | 448/7010<br>(0.0601) | - | - | - | NA | - | - | - |
| COI_BlocF | NA | 7126/3426<br>(0.6753) | 9257/1669<br>(0.8472) | 1809/303<br>(0.8565) | NA | NA | NA | 959/4955<br>(0.1622) | - | - | - | - |
| COI_BlocR | NA | 7428/5660<br>(0.56754) | NA | 1751/1450<br>(0.5470) | NA | NA | 406/5975<br>(0.0636) | NA | - | - | - | - |
| COI_BlocF_2 | - | - | - | - | NA | 404/2028<br>(0.1783) | 631/772<br>(0.4498) | 618/332<br>(0.6505) | NA | 779/2676<br>(0.2255) | NA | - |
| HCO_DPO | - | - | - | - | NA | 407/0<br>(1) | 156/0<br>(1) | NA | 445/144<br>(0.7555) | 463/91<br>(0.8357) | 175/0<br>(1) | 18488/196<br>(0.9895) |

## Results

### Distribution of *Rickettsia* in amphipod hosts in New Zealand

*Rickettsia* was detected in 26 of 69 locations (37.7%) from 3 families of amphipods: *Paracalliope* species complex (24/59 populations; 40.7%), *Paraleptamphopus* sp. (3/5 populations; 60%), and one undescribed family of Senticaudata (1/1 population; 100%) (Figure 1 and Supplementary Table 1). Because pooled samples were used, accurate prevalence in each population could not be obtained. However, a relative comparison was possible among the populations in which the same number of individuals per sample and the same total number was used (i.e. populations with a total of 48 individuals, with 12 samples each containing 4 individuals) (Supplementary Table 1). With parsimonious interpretation, among 18 populations, 12 populations showed at least 10% prevalence (>5 positive tubes/12 tubes tested). Seven populations showed at least 20% prevalence (>10 positive tubes/12 tubes tested). And three populations showed 100% positive tubes (12/12), with prevalence thus possibly ranging from 25-100%. Although *Rickettsia* was detected in both the North and South Islands, its distribution was confined to the southern parts of both islands (Figure 1).

### Genetic characterization of *Rickettsia* sequences

At least one *16S rRNA*, one *gltA*, or one *atpA* sequence was obtained from each of the population/species that were positive in the initial molecular screening (Supplementary Table 1). Specifically, 24 sequences of *16S rRNA*, 14 sequences of *gltA*, and 19 sequences of *atpA* were obtained. Also, 8 sequences of COI were added to our dataset. Fourteen genotypes were identified using *16S rRNA* sequences. All 16S sequences showed higher similarity to each other (>99.4%). All *gltA*, *atpA*, and COI sequences of Torix *Rickettsia* from amphipods showed high similarity to each other: >95%, >94%, >95%, respectively.

### Compiling molecular data on Torix *Rickettsia* from GenBank

A total of 183 nucleotide sequences of Torix *Rickettsia* were obtained from GenBank (Supplementary Tables 2-4). Specifically, 51 *16S rRNA* sequences were available from Amoeba, Annelida, Arachnida, Coleoptera, Diptera, Hemiptera, Hymenoptera, Psocoptera, Megaloptera, and an environmental sample representing 18 studies. A total of 68 sequences of *gltA* were obtained from Arachnida, Coleoptera, Diptera, Hemiptera, Hymenoptera, and Siphonaptera representing 12 studies. A total of 64 COI sequences were available from Amphipoda, Arachnida, Coleoptera, Diptera, Hemiptera, Hymenoptera, Megaloptera, and Odonata, representing 17 studies. Among these COI sequences, 42 sequences from 11 studies assigned rickettsial COI sequences to their invertebrate hosts. Since the very first misassignment in 2013, these mislabeled sequences have been deposited every year. Eight of these studies (representing 26 sequences) were involved with DNA barcoding and therefore voucher specimens (Supplementary Table 4).

### Phylogeny of *Rickettsia*

All three trees inferred by *16S rRNA*, *gltA*, and COI sequences clearly show two lineages within the genus *Rickettsia*: one clade of Torix *Rickettsia* and the other clade including all other 12 recognized groups within *Rickettsia* (Figures 3–5). A Bayesian tree based on *16S rRNA* sequences (Figure 3) shows that all sequences obtained from New Zealand amphipods belong to the Torix group of *Rickettsia*. Even when the *16S rRNA* conserved marker was used, all sequences obtained from amphipods (except S39_1542), were grouped in the same clade and distinct from other sequences, although this clade was not strongly supported (PP=0.84). Several subgroups were identified in the *gltA* tree (Figure 4). Most *Rickettsia* from amphipods were grouped within the same clade, similar to that revealed in the *16S rRNA* tree. Also, a Bayesian tree inferred from COI sequences (Figure 5) shows that *Rickettsia* from amphipods and some insects obtained in New Zealand are closely related, and the clade containing them is strongly supported (PP=0.96).

**Figure 3.**
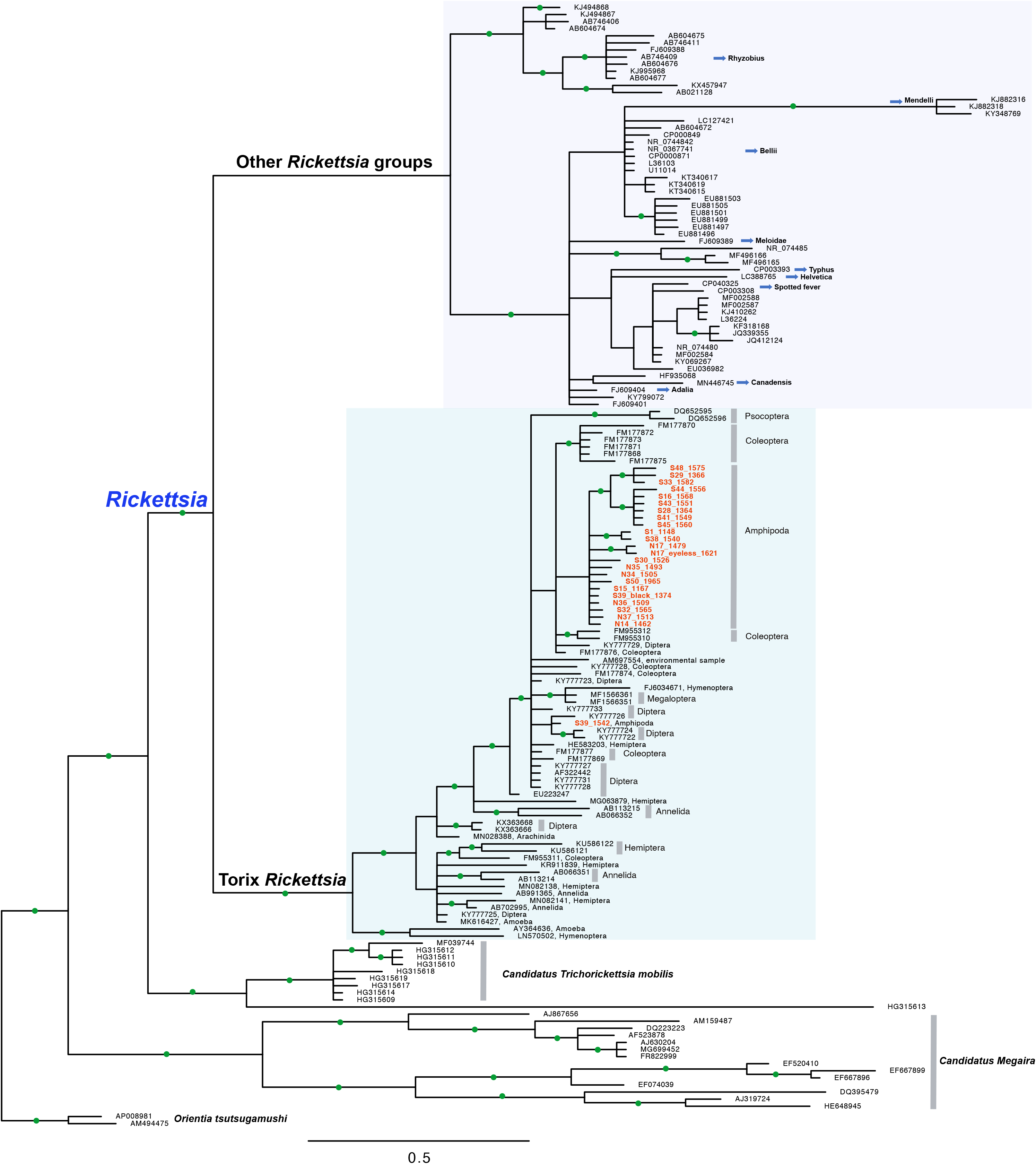
Bayesian tree of the genus *Rickettsia* based on 16S rRNA sequences. An alignment of 1,198 bp of 158 taxa was used. Two well-supported clades are shown within the genus *Rickettsia*. One is the Torix group which includes endosymbionts of diverse hosts (host taxa indicated on the right), and the other clade includes all other 12 recognized groups of *Rickettsia*. Nodes with a posterior probability higher than 0.9 are shown with green circles. Sequences obtained in this study are highlighted in orange colour.

**Figure 4.**
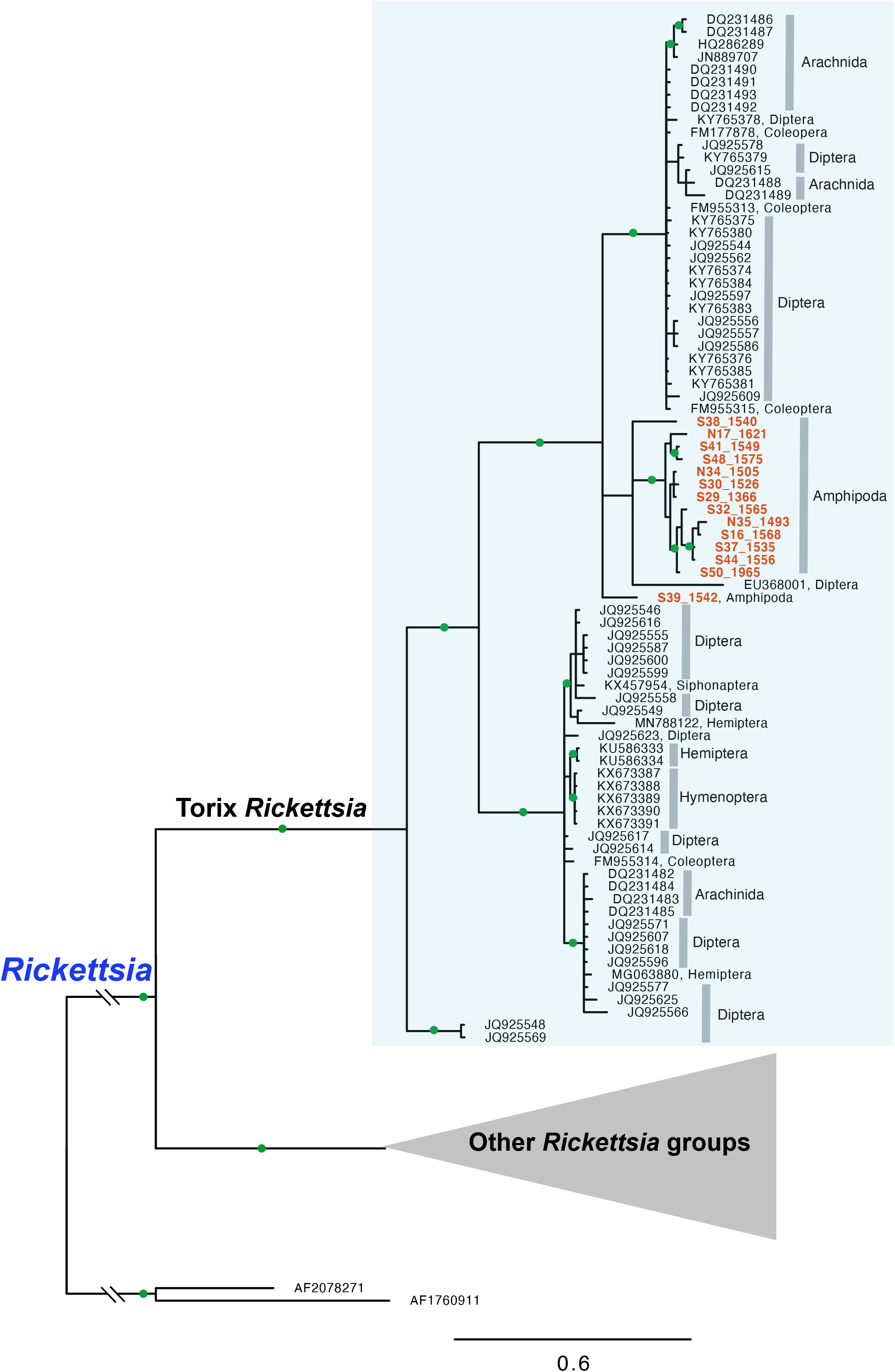
Bayesian tree of the genus *Rickettsia* based on an alignment of 765 bp of *gltA* sequences of 130 taxa (host taxa indicated on the right). Nodes with a posterior probability higher than 0.9 are shown with green circles. Sequences obtained in this study are highlighted in orange colour.

**Figure 5.**
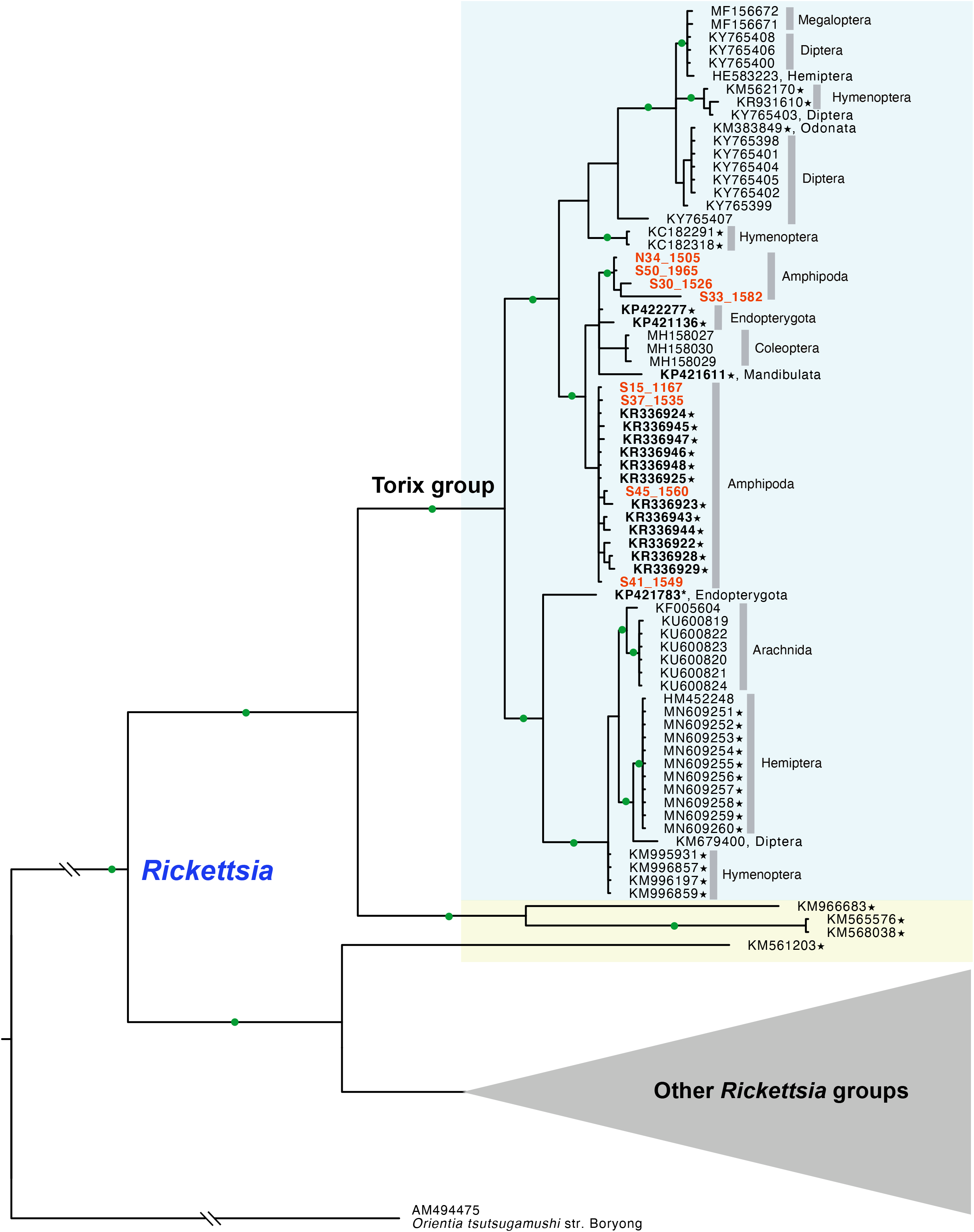
Bayesian tree of the genus *Rickettsia* based on an alignment of 559 bp of COI sequences of 114 taxa (host taxa indicated on the right). Nodes with a posterior probability higher than 0.9 are shown with green circles. Sequences that were misidentified as COI from invertebrate hosts, and initially not as rickettsial endosymbionts, in GenBank are highlighted with ★. Sequences obtained in this study are highlighted in orange colour. Sequences obtained in New Zealand are highlighted with bold text. Sequence similarity among Torix *Rickettsia* in the blue box is above 89%. Sequences that are similar to the other Torix *Rickettsia* but with lower similarity (80~82%), are highlighted within the yellow box.

### Testing and validating blocking primers

The ratio of the amplicons of host COI to *Rickettsia* COI (based on their expected fragment sizes) was calculated to compare the efficiency of each primer under different conditions (Table 2). Not all fragment analyses were successful, but we were able to compare some effects. Except for Bloc_R, which always resulted in amplifying an excess of *Rickettsia* COI (ratio 0.06~0.57), all primers showed some blocking effects. Specifically, Bloc_F, Bloc_F2, and HCO_DPO primers showed increased blocking effects (higher ratio of hostCOI:RickettsiaCOI) when more blocking primers were added. An increased number of PCR cycles resulted in a high number of host COI fragments. However, this increased somewhat the amplification of *Rickettsia* COI as well. Overall, HCO_DPO showed the highest efficiency among tested primers by preventing the amplification of *Rickettsia* COI, even at low concentrations.

## Discussion

Our nationwide molecular screening results show that the Torix group of *Rickettsia* is widespread in freshwater amphipod hosts in New Zealand, and is to our knowledge the first report of *Rickettsia* in crustacean hosts. Because of the lack of information on *Rickettsia* infections in other groups of amphipods in other parts of the world, when and how these bacteria colonised and spread among New Zealand amphipods remain in question. Because freshwater amphipods have limited dispersal abilities (Myers, 1993), the widespread distribution of Torix *Rickettsia* in New Zealand amphipods may be explained by an ancient acquisition followed by vertical transmission, or by many independent events of recent horizontal transmission from other organisms. Several lines of evidence indicate that both horizontal and vertical transmission may have played roles in spreading and maintaining these bacteria in New Zealand amphipod hosts. The monophyletic relationships of most *Rickettsia* from New Zealand amphipod populations inferred by *16S rRNA* and *gltA* sequences (Figures 3 and 4) seem to support the ancient acquisition scenario. Also, genetically closely related *Paracalliope* populations harbored the same genotype of *Rickettsia*, which also strongly supports their long-lasting relationship probably maintained by vertical transmission. Meanwhile, sharing of *Rickettsia* genotypes between sympatric amphipod species of different families suggests host shifts among genetically distant host species within the same order. Such a complex evolutionary history involving both vertical transmission and horizontal transfers has been reported for other insect/endosymbiotic systems (Duron et al., 2014). The Bayesian tree obtained with COI sequences provides some hints for horizontal transmission among amphipods and other arthropods (Figure 5). *Rickettsia* sequences from darkling beetles (*Pimelia* sp.) obtained in Europe were highly similar to the sequences identified in New Zealand amphipods (96~98% similarity) (López et al., 2018). Moreover, *Rickettsia* sequences obtained from several unspecified arthropod species (Mandibulata sp., Endopterygota sp., and Formicidae sp.) in New Zealand (although these were originally identified as invertebrate COI sequences) (Drummond et al., 2015) are closely related to those of New Zealand amphipods (96~99% similarity), providing strong evidence of recent horizontal transmission among them. Unfortunately, details regarding the host specimens and local origins of these ‘insect’ sequences in New Zealand are not available. Direct detection of *Rickettsia* from these arthropod species and multi-gene analyses will be necessary to elucidate their transmission routes.

Our findings support the early observation that the Torix group of *Rickettsia* may be highly associated with aquatic and damp environments (Weinert et al., 2009), which was based on the detection of this group in leeches, amoeba, Diptera, and Coleoptera (Dyková et al., 2003; Kikuchi et al., 2002; Küchler et al., 2009; Perlman et al., 2006). Pilgrim et al. (2017) provided support for this view by detecting *Rickettsia* from 38 % of *Culicoides* species tested and hypothesized that Torix *Rickettsia* may be dominant in insects with aquatic larval stages (Pilgrim et al., 2017). Horizontal transmission of Torix *Rickettsia* among genetically distantly related but spatially co-occurring species may have occurred frequently(Weinert et al., 2009). The high prevalence of Torix *Rickettsia* and their stable association with their hosts suggest negligible pathogenic effects of this group (Dyková et al., 2003; Kikuchi et al., 2002; Küchler et al., 2009; Wang et al., 2020). Some Torix *Rickettsia* may even be beneficial for their hosts. For example, infected leeches can have larger body sizes than uninfected individuals, although the possibility that larger individuals are more likely to acquire *Rickettsia* via horizontal transmission cannot be ruled out (Kikuchi and Fukatsu, 2005; Perlman et al., 2006). Ecological impacts of Torix *Rickettsia* on their hosts, and direct evidence of horizontal transmission among aquatic host groups, could be better answered with targeted community-level studies.

With the advancement of molecular techniques, our knowledge of the diversity of the sister groups of *Rickettsia* is also increasing and changing rapidly. Earlier studies focused mainly on the pathogenic and medically important species in arthropod hosts (Azad and Beard, 1998; Raoult et al., 2001). Until 2005, only two genera, *Rickettsia* and *Orientia*, were known within the family *Rickettsiaceae*, which now contains seven more genus-level taxa (Castelli et al., 2016; Sabaneyeva et al., 2018). All these new genera are exclusively found in aquatic environments, mostly within ciliate hosts. It seems that adaptation to the use of arthropod hosts occurred several times independently within the family *Rickettsiaceae* (Castelli et al., 2016). In addition, the phylogenetic status and relationships among *Rickettsia* groups are also changing. The Hydra group, which was once considered to be an ancient group within the genus *Rickettsia*, along with the Torix and Belli groups, is now regarded as a separate genus: *Candidatus Megaira* (Schrallhammer et al., 2013). Another recently recognized group, *Candidatus Trichorickettsia*, is now believed to be a sister clade to the genus *Rickettsia* (Sabaneyeva et al., 2018).

The Torix group is largely different from the other groups of *Rickettsia* in many respects, including host range and habitat. The Torix group includes not only endosymbionts of diverse aquatic invertebrates (that are more complex than ciliates), but also diverse terrestrial arthropod hosts. Also, the Torix group is genetically distinct from other groups of *Rickettsia*, which all are sister to Torix *Rickettsia*. Specifically, the genetic divergences between the Torix and the Bellii group are 96% in *16S rRNA*, 78% in *gltA*, and 76% in COI sequences. The delimitation criteria we used for the Torix group in this study were 98.1% in *16S rRNA*, 87.6% in *gltA*, and 89% in COI (broadly 80%; see Figure 5). Two genome sequences of Torix *Rickettsia* recently became available (Pilgrim et al., 2017; Wang et al., 2020). These genomes have the typical characteristics of *Rickettsia* (e.g. reduced genome size, and biosynthetic and catabolic capacity) but also have unique characteristics different from the other groups of *Rickettsia* (e.g. the presence of non-oxidative PPP, Methionine salvage pathway, and glycolysis). It would be interesting to see how these two sister lineages, one mainly pathogenic and the other nonpathogenic, evolved and diverged from their common ancestor.

Interestingly, COI sequences revealed a diversity of Torix *Rickettsia*, as much as other popular markers for Rickettsiales such as *16S rRNA* and *gltA*, even though COI has rarely been used as a marker of choice. Among 17 studies that have generated COI sequences of *Rickettsia*, only 3 were specifically intended to obtain COI sequences from *Rickettsia* (Gerth et al., 2017; MacHtelinckx et al., 2012; Pilgrim et al., 2017). The remaining 14 studies obtained *Rickettsia* COI sequences as a byproduct of other research objectives (i.e. host identification, population genetic studies, or DNA barcoding) with PCR using universal primers. *16S rRNA* is a widely used marker but may be too conserved to resolve phylogenetic relationships among closely related species, while the *gltA* gene shows more variability. Only 6 studies (including the present one) produced both *16S rRNA* and *gltA* sequences (Küchler et al., 2009; Noda et al., 2012; Pilgrim et al., 2017; Reeves et al., 2008; Wang et al., 2020). *gltA* sequences are not available for most Torix *Rickettsia* including endosymbionts of leeches and amoeba. Conversely, only *gltA* sequences are available for some species found in spiders and dipterans, which made it difficult to resolve the phylogenetic position of these rickettsial endosymbionts along with other *Rickettsia* (Goodacre et al., 2006; Perlman et al., 2006). ‘Limoniae’ and ‘Leech’ groups were used within Torix *Rickettsia* in some studies based on the *gltA* gene and concatenated sequences of *gltA* and *16S* rRNA genes, although the ‘Leech’ group was found not to be monophyletic (Gualtieri et al., 2017; Küchler et al., 2009). Our *gltA* tree showed two main lineages which correspond to the clades found in previous studies. Similarly, two main linages were identified in the Bayesian tree inferred from COI sequences. However, whether the clade containing all endosymbionts from spiders represents the ‘Leech’ group could not be confirmed without direct multi-gene data from the same group. It is likely that there are many more subgroups, given the limited number of targeted studies available to date.

Endosymbionts and vertically transmitted intracellular parasites are common in arthropod hosts (Rousset et al., 1992; White et al., 2013). In the context of the growing recognition of the ‘Holobiont’ concept (Minard et al., 2013; Thompson et al., 2014), obtaining bacterial sequences from DNA extracts from host tissue is not surprising. Most bacterial ‘contaminations’ are filtered out during processing of metabarcoding data (Leray et al., 2013; Siddall et al., 2009). The frequent recovery of COI sequences from Torix *Rickettsia* can be partly explained by their nucleotide sequence similarity with mitochondria. The Proto-mitochondrion (the hypothetical common ancestor of all mitochondria) is often recovered as a sister to the Rickettsiales or within the Rickettsiales (Roger et al., 2017). The alignment of COI sequences among several lineages of *Rickettsia* shows the high similarity between priming sites and the sequences used for universal primers (Figure 2). The priming site for the forward primer is 80% (20/25 nucleotides) identical to the LCO1490 sequences, and the priming site for the reverse primer is 84.6% (22/26 nucleotide) identical to the HCO2198 sequences. In addition, priming sites for universal primers are not conserved in many groups, which necessitates the need for group-specific or degenerate markers (Geller et al., 2013; Ward et al., 2005). However, this does not explain the more frequent reports of Torix COI in GenBank, because priming sites are also highly conserved in other groups of *Rickettsia*(Figure 2). Several explanations can be proposed. Other groups of *Rickettsia* often induce severe pathology and may therefore be rare in sampled populations. In contrast, the Torix group is generally non-pathogenic and often found at high prevalence in host populations. Therefore, several individuals from a given population may all be infected and could yield rickettsial COI, as illustrated in some previous studies (Ceccarelli et al., 2016; Lagrue et al., 2016).

These problems can be managed, as they are with *Wolbachia* (Smith et al., 2012). As mentioned earlier, Torix COI is highly conserved across diverse hosts. Therefore, comparing newly obtained (and suspicious) COI sequences with known Torix *Rickettsia* COI sequences can be easily done to distinguish Torix *Rickettsia*. Comparing sequences from the same taxon or genetically closely related groups could be useful. Checking for the presence of stop codons could largely decrease this problem, as for numts (Song et al., 2008). Bacterial sequences will show stop codons with the translation table for invertebrates, but will be in an open reading frame with the translation table for Bacteria. In addition to the high prevalence of Torix *Rickettsia* in many populations, high copy numbers of *Rickettsia* in host cells also make it difficult to obtain genuine host COI sequences, once a population is infected. In this case, applying blocking primers is a practical solution. Unfortunately, using blocking primers for *Rickettsia* does not always guarantee the amplification of host COI because other symbionts or parasites might still be amplified. Nevertheless, blocking primers can be widely used for any host groups that are infected by *Rickettsia*, and for both next-generation-sequencing as well as Sanger sequencing. These sequences should not be confounded with those of hosts, yet these ‘unwanted sequences’ or ‘contaminations’ can provide useful information about their endosymbionts and parasites. For example, the COI sequence obtained from a damselfly (GenBank ID: KM383849) suggests the presence of *Rickettsia* in this host group (order Odonata), in which Torix *Rickettsia* have never been reported. Targeted studies are likely to uncover a huge but under-detected diversity of Torix *Rickettsia*, and with more data, we will be able to answer questions regarding transmission, host switching, and the evolution of pathogenicity. Furthermore, detailed research on a finer scale is needed to elucidate the impact of these widespread endosymbionts on their diverse hosts.

## Acknowledgement

The authors gratefully thank Sophie Courjal, Eleanor Wainwright, Colin Décout, Brandon Ruehle, Jean-François Doherty, and Heloise Pavanato for their assistance in the field and/or laboratory. We appreciate Dr. Jim Lowry’s help for identifying amphipods. Also, we thank Dr. Tania King for introducing blocking primers and all practical help in the laboratory. We appreciate helpful comments provided by Dr. Pablo Tortosa and Jean-François Doherty. This research was supported by a University of Otago Doctoral Scholarship to EP.

## Author contributions

### Data availability statement

All sequences generated in this study were deposited in GenBank (Accession ID: MT507651-MT507674; MT515460-MT515486; MT524989-MT525002)

